# The iMab antibody selectively binds to intramolecular and intermolecular i-motif structures

**DOI:** 10.1101/2024.06.22.600195

**Authors:** Emanuela Ruggiero, Maja Marusic, Irene Zanin, Cristian David Peña Martinez, Janez Plavec, Daniel Christ, Sara N. Richter

**Affiliations:** Department of Molecular Medicine, University of Padua, 35121 Padua, Italy; Slovenian NMR Centre, National Institute of Chemistry, Hajdrihova 19, SI-1000 Ljubljana, Slovenia; Garvan Institute of Medical Research, Darlinghurst, Sydney NSW 2010, Australia; St Vincent’s Clinical School, Faculty of Medicine, University of New South Wales, Kensington, Sydney NSW 2010, Australia; Microbiology and Virology Unit, Padua University Hospital, 35121 Padua, Italy

**Author notes:** Senior authors (Janez Plavec, Daniel Christ, Sara N. Richter).

## Abstract

i-Motifs are quadruplex nucleic acid conformations that form in cytosine-rich regions. Because of their acidic pH dependence, iMs were thought to form only *in vitro*. The recent development of an iM-selective antibody, iMab, has allowed iM detection in cells, which revealed their presence at gene promoters and their cell cycle dependence. However, recently evidence emerged which seemed to suggest that iMab recognizes C-rich sequences regardless of their iM conformation. To further investigate the selectivity of iMab, we examined the binding of iMab to C-rich sequences, using a combination of pull-down and Western blot assays. Here we observe that the composition of buffers used during binding and washing steps strongly influences the selectivity of antibody binding. In addition, we demonstrate by NMR that several of the previously reported C-rich sequences, which were not expected to form iMs, actually form *intermolecular* iMs which are selectively recognized by iMab. Our results highlight the specificity of the iMab antibody, emphasize the importance of optimizing DNA concentrations, blocking and washing conditions, and confirm iMab selectivity not only for intramolecular iMs, but also for intermolecular iMs.

## INTRODUCTION

The high dynamics of DNA during cellular processes allows it to adopt several conformations alternative to the double helix, including quadruplexes such as i-motifs (iMs) and G-quadruplexes (G4s). iMs form within cytosine (C)-rich regions through the intercalation of hemi-protonated C^+^-C base pairs, when at least four tracts of at least two consecutive Cs are present (1). G4s occur within guanine (G)-rich regions when two or more G-tetrads, which are associations of four Gs, self-stack (2). While G4s have been extensively studied thanks to the availability of specific antibodies (3) and ligands that recognize and bind them (4, 5), iMs have been much less explored. Prediction algorithms have located putative iM-forming sequences in key regulatory regions, such as gene promoters, centromeres and telomeres (6). Many studies have investigated iM formation experimentally *in vitro*, contributing to the understanding of its folding factors (6). The development of the first anti-iM antibody, iMab (7), was a breakthrough in iM research, leading to the detection and mapping of iMs in cells (8). iMab has been successfully used in several *in vitro* techniques, such as dot blot (9, 10) and pull-down (11) assays, and has been used to develop a custom microarray for the screening of thousands of iM-forming sequences (12), which has expanded iM structural characterization. In cells, immunofluorescence with iMab localized iMs in the nucleus of several human cell lines (7), and showed that their number was cell cycle dependent. iMs were reported to be most abundant during active transcription phases, such as G1, and to decrease in subsequent phases (13), implicating iMs in cell regulatory roles. Recently, iMab has been used in different high-throughput sequencing-based techniques, iMab-IP-Seq (9, 14) and CUT&Tag (11), providing precise information on the distribution and location of iMs in the whole genome. In the first case, the analysis was performed on purified DNA extracted from the rice plant, where iMs showed an intrinsic subgenomic distribution and *cis*-regulatory function (9); methylation of the immunoprecipitated region was found to strongly influence iM formation, revealing new aspects in genome regulation (10). In the second case, the analysis was performed on purified human genomic DNA, demonstrating the wide distribution of sequences capable of iM formation, which are common among highly expressed genes and those upregulated in G0/G1 cell cycle phase (14). In the context of chromatin, our group recently applied iMab to the CUT&Tag protocol on two different human cells. We found that iMs in cells are mainly located at actively transcribing gene promoters in open chromatin regions, and that their abundance and distribution are specific to each cell type. iMs with both long and short C-tracts were recovered, and their folding was further confirmed *in vitro* (11). The data obtained with the iMab-CUT&Tag were then used to develop a machine learning pipeline called iM-Seeker, which aims to predict both the folding status and the structural stability of iMs, based on experimental evidence (15).

In contrast to the above studies, a recent NMR study indicated that synthetic iM-forming oligonucleotides inserted into the nucleus are often not folded, suggesting that the nuclear environment may not be broadly supportive of iM folding (16). Recently, a preprint by Boissieras et al. suggested that the iMab antibody is capable of binding to C-rich synthetic DNA oligonucleotides independent of iM formation (17). These apparent discrepancies are not unexpected due to the transient nature of iM formation, and the use of different experimental methods and conditions, including buffer conditions and DNA organization (purified genomic DNA vs chromatin vs synthetic oligonucleotides) (18). To address these issues, we have investigated the selectivity of iMab towards C-rich sequences using a pull-down/western blot (WB) approach (11). We report the importance of optimizing condition of each key step, highlight the specificity of iMab, and that this antibody selectively recognizes both intramolecular and intermolecular iMs.

## MATERIALS AND METHODS

### Oligonucleotides

Oligonucleotides used in the CD and pull-down assays were purchased from Sigma-Aldrich and are listed in Table 1. For the pull-down assay, a biotin-TEG residue was added at the oligonucleotide 3’-end.

**Table 1.**
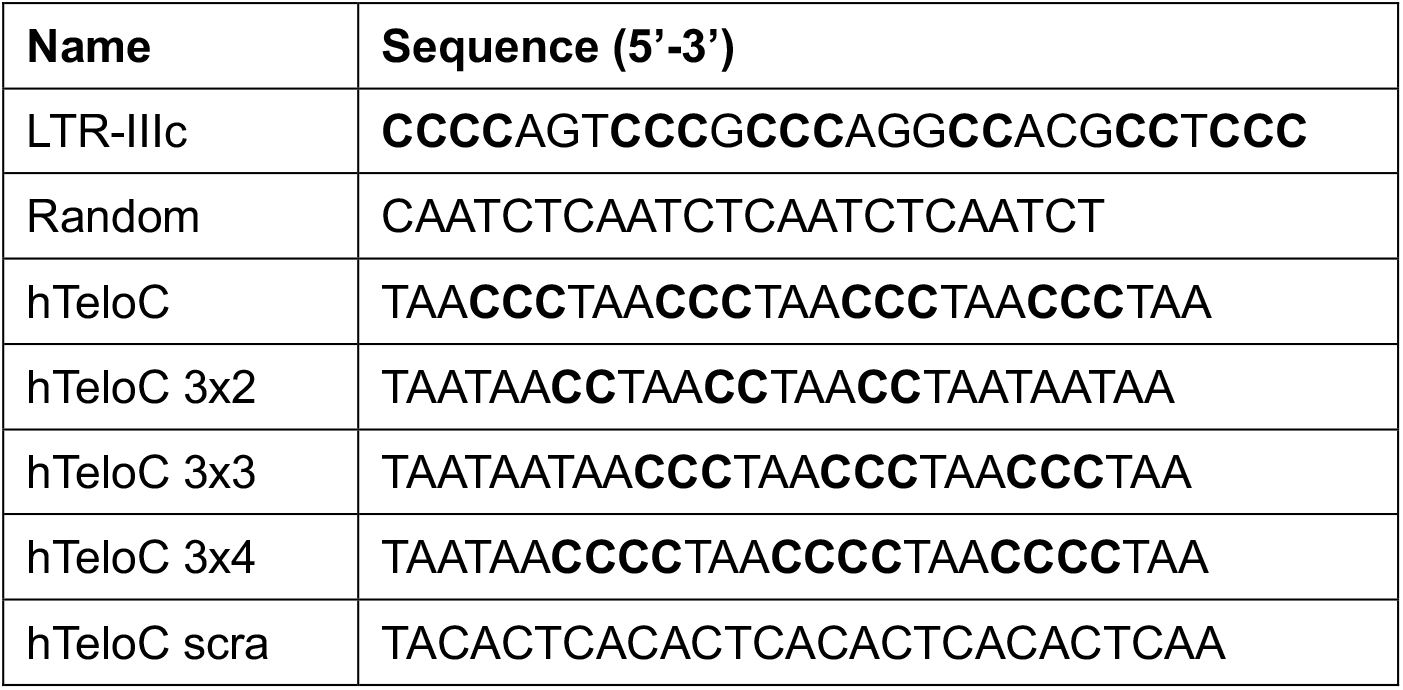
Oligonucleotides used in this study. C-tracts composed of two or more Cs are shown in bold.

### Circular Dichroism

Oligonucleotides used for circular dichroism (CD) analysis were diluted to 3 μM in 20 mM phosphate buffer, 80 mM KCl at pH 5.4, 6, and 7.4. Samples were heated at 95°C for 5 min and then slowly cooled to room temperature overnight. 5 mm and 1 mm optical-length quartz cells were used to record CD spectra on a Chirascan-Plus equipped with a Peltier temperature controller. CD spectra were performed at 20°C and data were acquired from wavelength 230 to 320 nm. CD data were baseline-corrected and the observed ellipticities were converted to mean residue ellipticity according to θ = degree x cm^2^ x dmol^-1^ (molar ellipticity). CD analysis was performed in duplicate and values were plotted using R as the mean of the two independent replicates (19).

### iMab Pull-down and Western-blot

Pull-down coupled with WB was adapted from previously described procedures (11). In brief, samples were prepared by diluting biotinylated oligonucleotides (Table 1) to 1.5 μM or 0.3 μM in 100 mM phosphate buffer pH 6, denaturing at 95°C for 5 min and cooling overnight. Upon immobilization of the samples on streptavidin-coated magnetic beads (Dynabeads™ M-280 Streptavidin, ThermoFisher Scientific, #11205D), oligonucleotides were incubated with 100 or 10 ng FLAG-tagged iMab (Absolute antibody, #Ab01462-30.135) for 1 h in ice bath in the appropriate binding buffer (Tris-HCl pH 7.5 10 mM, MgCl_2_ 1 mM, KCl 10 mM, DTT 1 mM), enriched with blocking agents and NaCl as described in Figure 2. Samples were washed four times in wash buffer (50 mM Tris-HCl pH 7.5/NaCl at different concentrations) and once with PBS. WB was performed according to known procedures (20). Samples were loaded on a 10% SDS-PAGE and the gel was then transferred on a PVDF membrane. The latter was blocked in 2.5% PBS-milk buffer for 1 h, incubated with the anti-FLAG antibody 1:1000 (Sigma Aldrich, #F3165), washed in 0.1% PBS-tween, and finally incubated with secondary goat anti-mouse 1:5000 HRP antibody (Merck-Millipore #12-349). Images were acquired on the Alliance Uvitec (Uvitec Ltd. Cambridge, Cambridge, United Kingdom) instrument by HRP bioluminescence measurement.

### Nuclear Magnetic Resonance

DNA oligonucleotides were synthesized with DNA/RNA H-8 K&A Laborgereate GbR synthesizer using standard solid-phase phosphoramidite chemistry in a DMT-ON mode, deprotected with AMA, purified with Glen-Pak™ cartridges and desalted on ÄKTA Purifier with HiPrep 26/10 Desalting column. Samples were prepared at 0.1 or 1 mM oligonucleotide concentration in 80 mM KCl, 20 mM potassium phosphate buffer at pH 5.4 or 6 and 10% D_2_O. Samples were annealed by heating to 95°C and cooling to room temperature overnight. 1D ^1^H NMR spectra were recorded on Bruker AVANCE Neo 600 MHz NMR spectrometer with 256 scans, spectral width of 24.5244 ppm and interscan delay of 2 s. Excitation sculpting was used for the suppression of water signal. Spectra were processed, analyzed, and integrated with TopSpin 4.1.4 (Bruker).

## RESULTS

### Optimization of iMab binding conditions

In order to expand our knowledge of iM structures formed by C-rich sequences, we set up a combined pull-down/WB experimental approach to assess iMab specificity. We selected seven sequences (Table 1) as follows: two previously characterized iMs, namely the LTR-IIIc sequence located within the HIV-1 virus promoter (21) and the hTeloC sequence from the C-rich region of the human telomere (22), both reported to fold into stable iMs, as positive control sequences; a Random sequence unable to fold into stable DNA secondary structures as negative control. In addition, following Boisseras et al. (17), we included three other sequences derived from the hTeloC, all containing three C-tracts of two (hTeloC 3×2), three (hTeloC 3×3) or four (hTeloC 3×4) Cs each. Finally, an additional negative control (hTeloC scra), derived from the reshuffling of the hTeloC primary sequence, was included.

First, we defined the folding profile of the selected sequences by circular dichroism (CD) analysis performed at increasing pH from 5.4 to 7.4 (Figure S1). We observed that the positive controls LTR-IIIc (Figure S1A) and hTeloC (Figure S1C) were iM folded in a pH-dependent manner, showing a positive peak around λ = 285 nm and a negative one around λ = 260 nm, as expected (23). The hTeloC scra sequence (Figure S1B) displayed a CD spectrum characteristic of unfolded DNA (24). For the hTeloC variants, hTeloC 3×4 was found to be folded at acidic pH (Figure S1F), while the hTeloC 3×2 showed an unfolded CD signature (Figure S1D). The hTeloC 3×3 sequence, which contains three CCC tracts, showed a shift of the major peak towards higher wavelengths, suggesting partial folding (Figure S1E). As the subsequent pull-down/WB analysis was performed on biotinylated sequences, we also evaluated the CD spectra of these sequences and observed no major changes (Figure 1), indicating that the presence of the terminal biotin residue does not detectably affect their three-dimensional fold.

**Figure 1.**
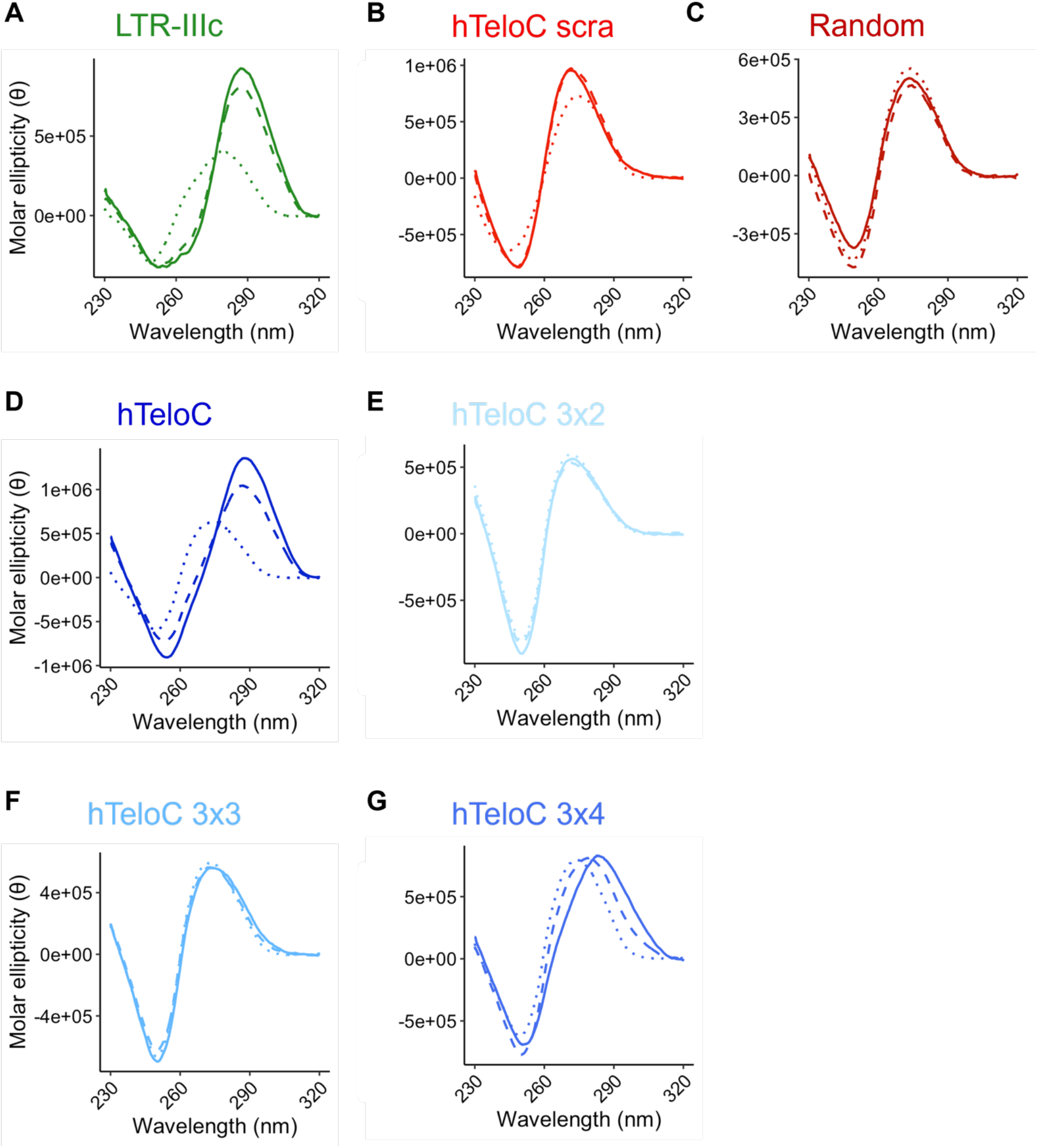
CD analysis of the biotinylated sequences used in the pull-down/WB assay. Samples were prepared in phosphate buffer at pH 5.4 (plain line), pH 6 (dashed line) and pH 7.4 (dotted line). θ = deg × cm2 × dmol-1.

In the pull-down assays, the folded biotinylated oligonucleotides are first immobilized onto streptavidin-coated magnetic beads and incubated with the protein of interest (in our case the iMab antibody). Beads are then washed several times to remove non-specific and weak interactions (Figure 2A); the retained bound antibody is visualized by WB. We initially selected a known positive iM (LTR-IIIc) (21) and a negative sequence (hTeloC scra) and started optimizing both washing and binding conditions to obtain the best selectivity settings for iMab binding. First, we maximized the efficiency of the washing buffer by testing the ionic strength, using NaCl concentrations from 0.3 M to 2.4 M (Figure 2B). We observed that, in general, LTR-IIIc showed a more intense band in the WB compared to the hTeloC scra, and, for both sequences, the binding became weaker with increasing NaCl concentrations. Therefore, in the following steps, we used concentrations of 0.15 or 0.3 M NaCl during the washing phase. As for the binding buffer, we evaluated the effect of the presence of different blocking agents, such as skim milk and BSA protein, at different concentrations as shown in Figure 2C. We also included in the analysis a sample containing additional NaCl, to increase the ionic strength during the binding step. We confirmed that iMab preferentially binds to the positive iM oligonucleotide, LTR-IIIc, and to a much lesser extent to the hTeloC scra. We observed that skim milk was more efficient than BSA in improving the selectivity for the iM folded sequence. The addition of ionic strength also went in the same direction. Therefore, we combined the results obtained in the first two settings and tested i) the concentration of skim milk (0.5% or 2.5%) to which 0.15 M NaCl was added; ii) the ionic strength of the washing buffer (0.15 M or 0.3 M) (Figure 2D). Overall, the most selective iMab binding was obtained with 2.5% skim milk and 0.15 M NaCl in the binding buffer and 0.15 M NaCl in the washing buffer. Based on these data, we applied the optimized protocol to all the selected sequences (Figure 2E, top panel) and observed that iMab was able to bind to all iM-forming oligonucleotides (LTR-IIIc, hTeloC and hTelo C 3×4), while the Random fragment showed no band. However, a clear band was observed for the hTeloC 3×2 and 3×3 sequences, and a weaker band was observed for the negative control hTeloC scra. We next evaluated the amount of antibody used in the analysis, by reducing iMab concentration by 10-fold (Figure 2E, bottom panel): we observed no reduction in band intensity for any of the tested sequences, except for hTeloC scra, for which binding largely disappeared.

**Figure 2.**
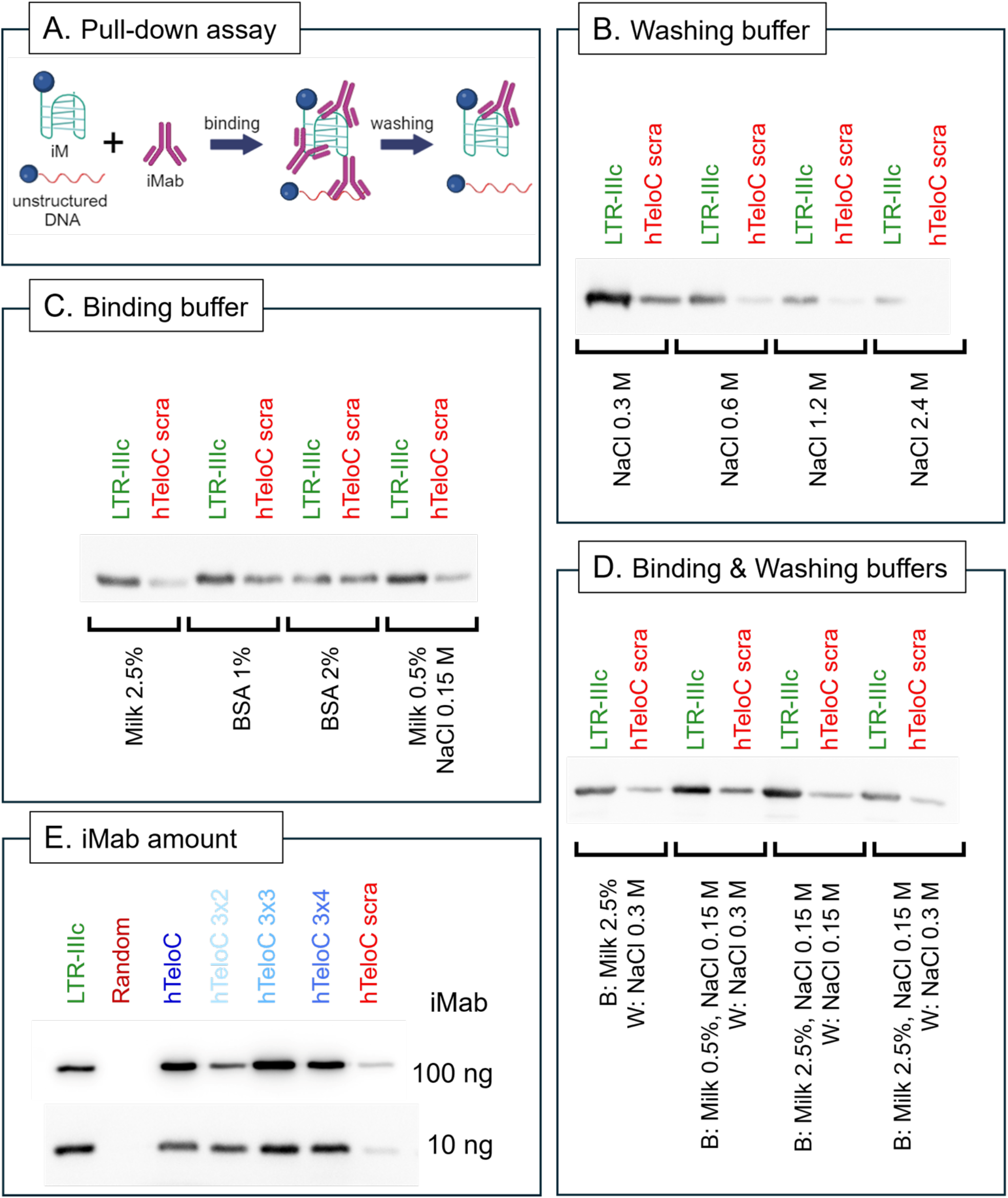
Assessing iMab selectivity by pull-down/WB assay. A) Cartoon model of the pull-down experimental approach. The biotinylated oligonucleotides were immobilized on streptavidin beads and incubated with iMab in the binding buffer indicated in panels C and D. Subsequently, the obtained complexes were subjected to multiple wash steps in the buffer indicated in panels A and D to remove weak and unspecific interaction as. B) Evaluation of the effect of increasing ionic strength in the washing buffer, from 0.3 M to 2.4 M NaCl. C) Evaluation of the effect of blocking agents (skim milk or BSA at different concentrations) and ionic strength (NaCl) in the binding buffer. D) Binding (B) and washing (W) buffer optimization. E) Pull-down followed by WB analysis on all sequences of interest at two different iMab concentrations, performed in the optimized conditions: binding buffer (milk 2.5%, NaCl 0.15 M), washing buffer (NaCl 0.15 M).

### iMab binds to both intramolecular and intermolecular iMs

Although we were able to identify the best conditions for iMab to recognize its target, we still observed a clear WB band for the hTeloC 3×2 and 3×3 sequences, which, based on their sequence, would not form intramolecular iM structures. However, we reasoned that both oligonucleotides could form intermolecular iM structures (25), which would explain their binding to iMab. Multimolecular iMs have mainly been studied for their application in nanotechnology and it is generally reported that they form in sequences with short C-tracts (25, 26), and therefore show lower stability than intramolecular structures (27). To prove our hypothesis, we performed ^1^H NMR analysis at two different oligonucleotide concentrations (Figure 3A), since higher DNA concentrations stimulate the formation of intermolecular structures (28). Signals between δ 15 and 16 ppm are characteristic of protonated C residues found in iMs (28) and were observed for hTeloC and hTeloC 3×4 oligonucleotides, regardless of DNA concentration and at both pH 5.4 and 6.0, indicating the formation of stable iMs. Interestingly, the hTeloC 3×2 and 3×3 sequences also displayed signals from protonated C residues, albeit predominantly at high DNA concentrations and with a greater dependence on pH, supporting the formation of intermolecular structures with low stability. To deepen our analysis, we examined the NMR melting profiles (Figure 3B), which showed a significant increase in stability for hTeloC, hTeloC 3×4 and hTeloC 3×3 oligonucleotides (8-11°C), but lower stabilization for hTeloC 3×2 (3°C), which was also the least stable iM. These results, along with the concentration-dependent shift in the melting temperature of all oligonucleotides (Figure 3B), are consistent with the formation of intermolecular iMs (27).

**Figure 3.**
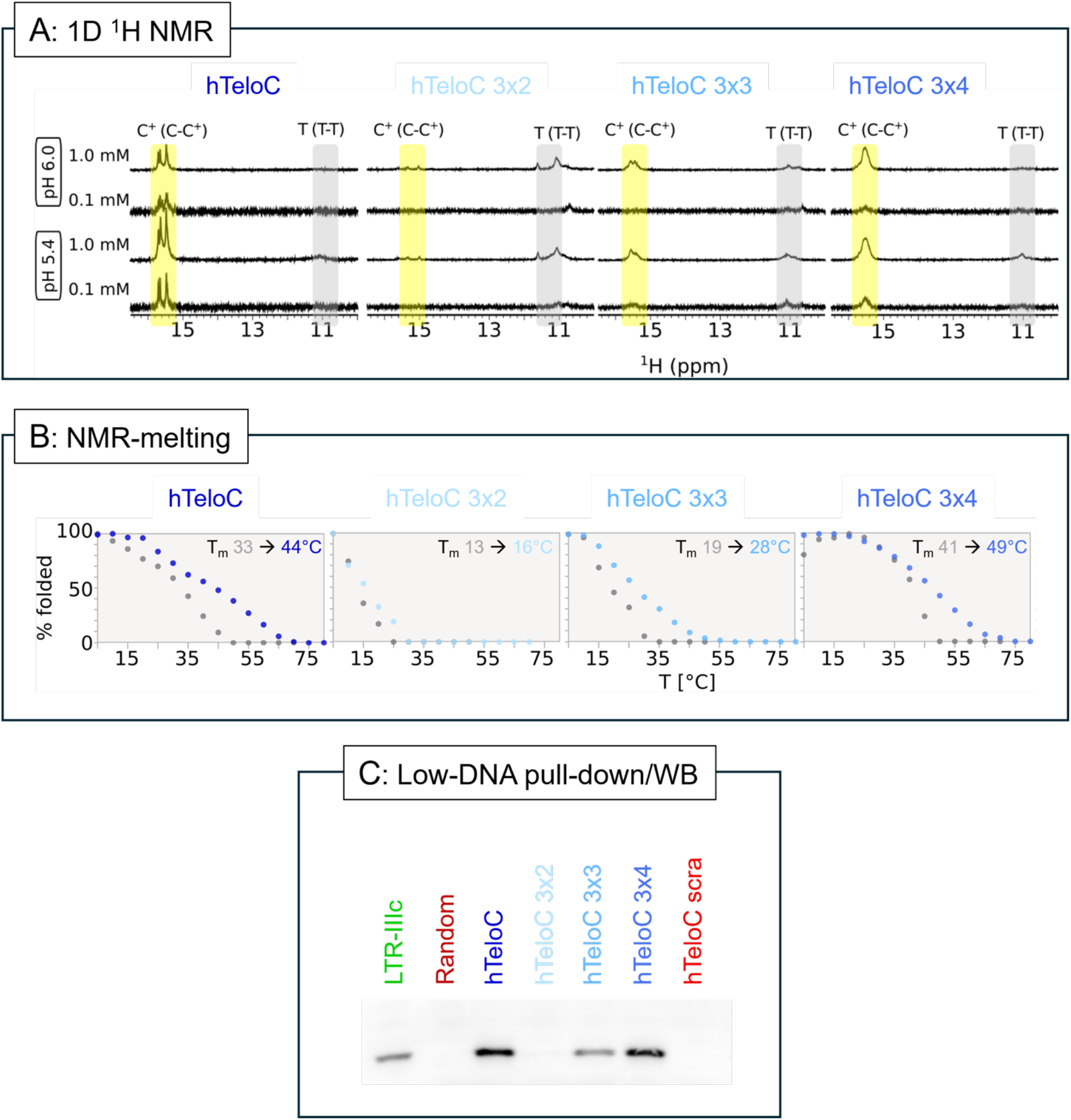
iMab binding profile to intermolecular iMs. A) Imino region of 1D ^1^H NMR spectra recorded at 5 °C, at different pH and at oligonucleotide concentrations. Regions characteristic for signals of protons included in non-canonical C-C^+^ and T-T base-pairs are highlighted by the yellow and grey areas, respectively. B) T_1/2_ at two different oligonucleotide concentrations obtained from NMR melting experiments based on the intensity of the signals in the imino region. Data points in the melting profiles displayed in gray represent data at 0.1 mM oligonucleotide concentration, while data points in shades of blue represent data at 1.0 mM oligonucleotide concentration. C) Pull-down/WB performed at low DNA concentration, 300 nM, with 10 ng iMab per sample.

We hence performed pull-down/WB analysis at lower DNA concentrations to minimize the formation of intermolecular iMs (Figure 3C). We found that iMab binding was robust for monomolecular iM-forming sequences, i.e. those sequences that showed NMR iM signature peaks at both high and low concentrations (LTR-IIIc, hTeloC, hTeloC 3×4), whereas it was strongly reduced (hTeloC 3×3) or completely lost (hTeloC 3×2) for those sequences that could only form intramolecular iMs, i.e. those sequences that showed NMR iM signature peaks only at high DNA concentrations. No band was observed in the negative controls. Overall, our data confirm the selective binding of iMab to iM-forming sequences and show for the first time that the antibody is able to recognize both intramolecular *and* intermolecular iM structures.

## DISCUSSION

The dynamic state of non-canonical nucleic acid structures has made their detection a challenge, especially within the environment of the cell. The development of the iMab antibody, the first to recognize iM DNA structures, has given a major boost to the research in this field. In fact, iMab has been used in several types of assays in recent years: different laboratories have shown that iMs are folded in the genome of different species (9, 11) and are localized in the nucleus of human cells (13). Recently, however, the selectivity of iMab has been challenged in a pre-print by Boissieras et al. who reported that iMab recognized C-rich sequences independently of iM-folding (17).

Prompted by the conflicting evidence, we set out to thoroughly investigate the selectivity of iMab using pull-down assays. We found that iMab binding is strongly influenced by experimental conditions and that the presence of the appropriate blocking agents and optimized salt concentrations are critical to avoid non-specific interactions. The use of skim milk and NaCl greatly improved the specific binding to iM forming sequences compared to unstructured DNA (Figure 2 B-D). However, it should be noted that the optimized conditions reported here for the pull-down/WB experimental approach may not be readily applicable to other techniques, and any new technique, such as immunofluorescence or immunoprecipitation, requires proper setup and critical analysis of the results. For example, Schneekloth et al. developed iMab-based microarrays, in which different pH levels and crowding conditions were shown to cause slight changes in the iMab binding pattern; however, non-iM-forming sequences did not give a measurable response, regardless of the settings, thus confirming iMab specificity under these specific conditions (12). Ma et al. have shown that iM-folded synthetic oligonucleotides and sonicated chromatin from cells provided similar binding in a iMab-dot blot assay, while negative controls showed no signals, confirming iMab selectivity under specific experimental conditions (9). iMab selectivity was further supported by whole genome sequencing approaches, where iMab-immunoprecipitated sequences were significantly enriched when compared to the negative control, both in a fragmented genomic DNA context (9, 14) and in unfixed chromatin (11). In unfixed chromatin, the results obtained with iMab were strikingly similar to those obtained with BG4 for G4s (11, 29, 30), with the main difference being the level of expression of the genes presenting folded iMs versus G4s in their promoter. If iMab had recognized all C-rich sequences, this would not have been the case. In addition, iMab peaks were found to be enriched in iM forming sequences, that were shown to fold *in vitro* at acidic pH. This last aspect further emphasizes that different folding conditions are expected *in vitro* and in cells.

An additional point of interest arising from our present results is the concentration of iMab and the target oligonucleotide in the pull-down assay. iMab binds its iM target with high (nanomolar) affinity (7) and consequently retained binding strength towards iMs even when strongly diluted (Figure 2E). It may therefore be advisable to use as little amount of iMab as possible for each experiment to avoid unnecessary saturation and to increase selectivity. Similarly, when titrating the target DNA concentration, it should be considered that high DNA concentrations can stimulate the formation of intermolecular quadruplex structures (25, 26). We have shown by NMR that the hTeloC variants used by Boissieras et al. and in this study are capable of forming intermolecular iMs (Figure 3A-B), providing a direct explanation for the apparent discrepancies reported by the authors, and confirming the specificity of the iMab antibody.

Our results provide new insights into the iM field and demonstrate that iMab can be used to detect intermolecular iMs *in vitro*. With the presence of intermolecular G4s in cells recently reported (31), this raises the intriguing possibility of such intermolecular structures could also be formed by iMs *in vivo*, and that such a scenario could be confirmed by iMab binding.

## AUTHOR CONTRIBUTION

E.R. performed the pull-down and WB experiments, analyzed the data, supervised the project, and wrote the paper; M.M.: performed the NMR experiments and analyzed the data; I.Z. performed the CD experiments and analyzed the data; J.P.: acquired funds and supervised the NMR project; C.P.M. and D.C. provided reagents, advised on conditions, and edited the paper; S.N.R. conceived and supervised the project, acquired funds, and wrote the paper.

## ACKNOWLEDGMENTS

We thank the BIONIC group at the University of Padua for the helpful discussion.

