## Supplementary Figure S1 for "The iMab antibody selectively binds to intramolecular and intermolecular i-motif structures"

[#] Senior authors

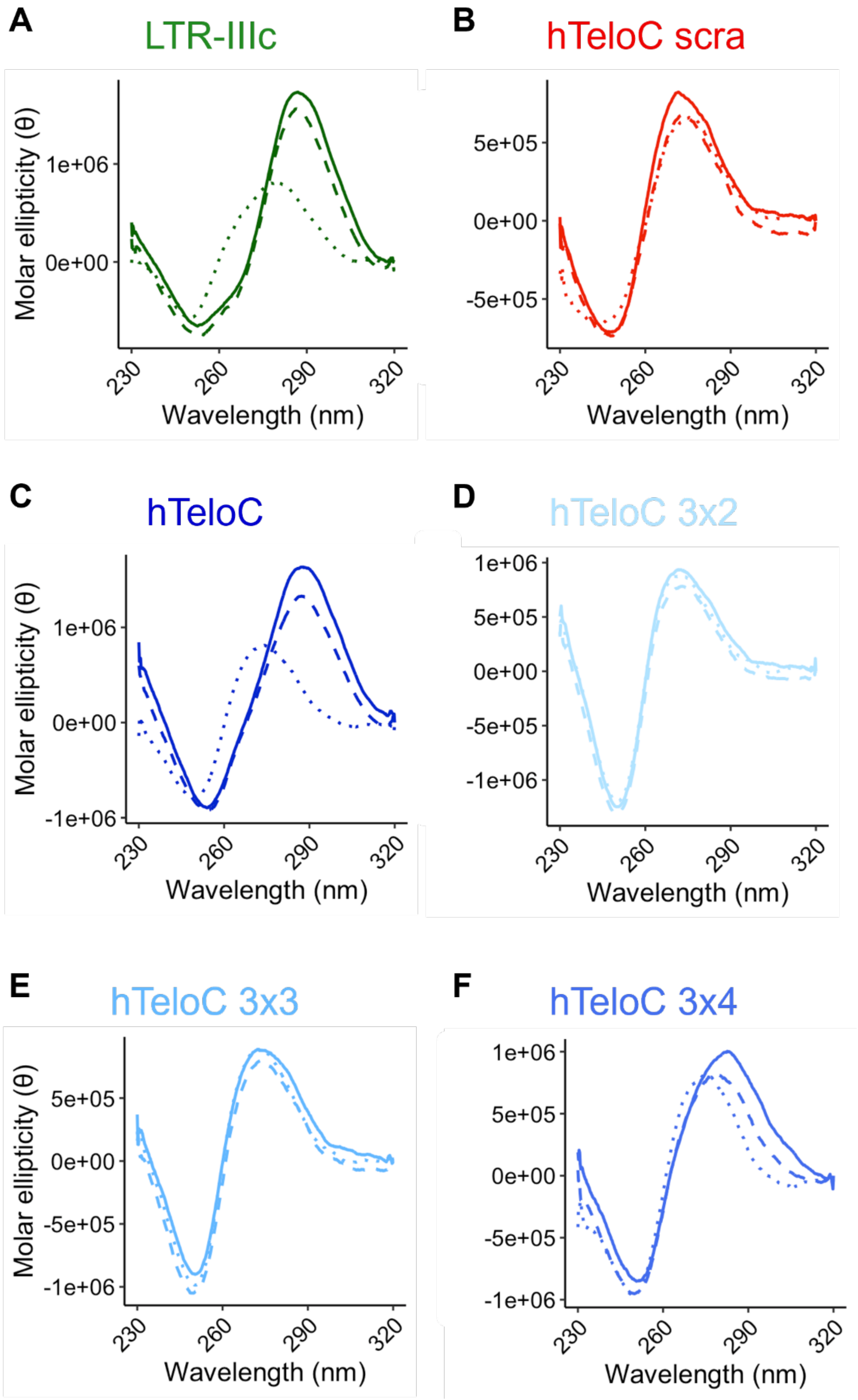


Figure S1. Circular dichroism analysis of selected sequences. Samples were prepared in phosphate buffer at pH 5.4 (plain line), pH 6 (dashed line) and pH 7.4 (dotted line). $\theta = \text{deg} \times \text{cm}^2 \times \text{dmol}^{-1}$.
